# A lineage-related reciprocal inhibition circuitry for sensory-motor action selection

**DOI:** 10.1101/100420

**Authors:** Benjamin Kottler, Vincenzo G. Fiore, Zoe N. Ludlow, Edgar Buhl, Gerald Vinatier, Richard Faville, Danielle C. Diaper, Alan Stepto, Jonah Dearlove, Yoshitsugu Adachi, Sheena Brown, Chenghao Chen, Daniel A. Solomon, Katherine E. White, Dickon M. Humphrey, Sean M. Buchanan, Stephan J. Sigrist, Keita Endo, Kei Ito, Benjamin de Bivort, Ralf Stanewsky, Raymond J. Dolan, Jean-Rene Martin, James J. L. Hodge, Nicholas J. Strausfeld, Frank Hirth

## Abstract

The insect central complex and vertebrate basal ganglia are forebrain centres involved in selection and maintenance of behavioural actions. However, little is known about the formation of the underlying circuits, or how they integrate sensory information for motor actions. Here, we show that paired embryonic neuroblasts generate central complex ring neurons that mediate sensory-motor transformation and action selection in Drosophila. Lineage analysis resolves four ring neuron subtypes, R1-R4, that form GABAergic inhibition circuitry among inhibitory sister cells. Genetic manipulations, together with functional imaging, demonstrate subtype-specific R neurons mediate the selection and maintenance of behavioural activity. A computational model substantiates genetic and behavioural observations suggesting that R neuron circuitry functions as salience detector using competitive inhibition to amplify, maintain or switch between activity states. The resultant gating mechanism translates facilitation, inhibition and disinhibition of behavioural activity as R neuron functions into selection of motor actions and their organisation into action sequences.

## INTRODUCTION

The selection and maintenance of a behavioural action is the outcome of specific neural interactions mediated by circuits that have evolved to coordinate adaptive behaviours. Essential to such circuits is their ability to organize sequences of actions by facilitating appropriate motor programmes while inhibiting competing ones. The underlying action selection mechanisms involve defined centres in the forebrain of arthropods and vertebrates including, respectively, the central complex (CX) and the basal ganglia (BG)^1^

In vertebrates, the BG mediate action selection by two main pathways that project from medium spiny GABAergic interneurons (MSN) of the striatum to the outer and inner shell of the Globus Pallidus. Striatal MSN that express dopamine D1 receptors form the direct pathway, whereas MSNs expressing dopamine D2 receptors constitute the indirect pathway. Although initially thought to act antagonistically, functional imaging studies reveal simultaneous activity of both direct and indirect pathways during behavioural sequences^2–4^, suggesting that both are involved, with different roles, in selecting and enhancing motor programs while inhibiting competing ones^5–10^. However, despite these advances, little is known about the underlying neural mechanisms, or how they are formed.

We recently identified extensive correspondences in neural organization and function of the BG with the CX of mandibulate arthropods^1,10^. In *Drosophila*, the CX comprises five centres: the protocerebral bridge (PB), fan-shaped body (FB), ellipsoid body (EB), noduli (NO) and lateral accessory lobes (LAL) (Figure 1). Similar to the BG, the CX, in particular its EB, plays an essential role in a multitude of behavioural manifestations, including goal-directed locomotion, spatial orientation, learning and memory, as well as a role in sleep, attention, arousal, and decision-making^11–27^. All of these behaviours require the selection and maintenance of motor actions, or their active suppression. Thus, within the context of the proposed homology of CX and BG circuitries, the many tools available for targeted cell and circuit-specific manipulations of the EB in *Drosophila*^28^, makes the invertebrate system ideal for investigating neural mechanisms and computations underlying action selection.

**Figure 1.**
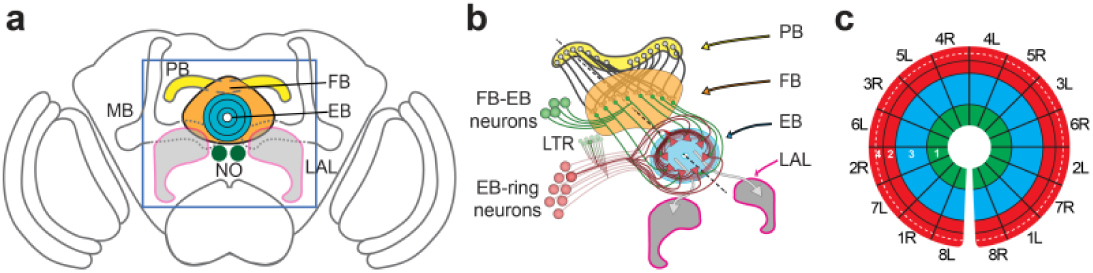
Structural organisation of the central complex and ellipsoid body. (**a**) Cartoon of adult *Drosophila* brain showing central complex ground pattern (PB, protocerebral bridge; FB, fan-shaped body; EB, ellipsoid body; NO, noduli; LAL, lateral accessory lobes - MB, mushroom bodies are shown for orientation); box indicates region in b. (**b**) CX neuropils are interconnected by projections from columnar neurons and tangential neurons (FB-EB neurons and EB-ring neurons). Histological, immunocytochemical, and clonal analyses demonstrate that columnar projection neurons connect all CX substructures and are subdivided into modules encoding spatial information about sensory events. In insects like grasshopper and *Drosophila*, 16 modules can be identified, which integrate sensory information that is processed in 8 PB units per hemisphere, representing the entire sensory space. Decussation from the PB to FB and EB leads to convergence of sensory representations, such that information from each hemisphere is integrated according to sensory segments. The CX receives sensory and somatosensory information from the protocerebrum via columnar projection and tangential neurons, but also via the lateral triangles (LTR), a microglomerular structure located between R neurons and EB neuropil. Integrated sensory and somatosensory associations received and processed in PB, FB and LTR, are weighted and selected in the EB. (c) In *Drosophila*, the EB forms a toroidal or ring-like neuropil that resembles a ‘closed arch’. Projections from tangential R neurons divide it into at least 4 different layers (white numbers 1-4). Sensory associations from PB and FB converge onto EB modules that can be subdivided according to PB-FB input spanning the left and right hemispheres.

Here we use genetic tracing from the embryo into the adult to show that all four subtypes of EB ring neurons (henceforth called R neurons, R1-4) in the adult *Drosophila* CX derive from embryonic neuroblast ppd5. Anatomical and genetic analyses reveal that R neurons establish a connectivity network among clonally related sister cells. Calcium-imaging and electrophysiological recordings demonstrate that R2-4 neurons are inhibitory GABAergic and are inhibited by GABA-A receptor signalling. We show that targeted manipulations of subtype-specific R neurons reveal that EB microcircuitry is involved in sensory-motor transformation required for the selection and maintenance of motor actions and their organisation into action sequences. A neural model of the EB R neuron circuit based on these structural and functional data suggests computational mechanisms mediating sensory-motor transformation and motor action selection in *Drosophila.*

## RESULTS

### R neurons derive from embryonic neuroblast ppd5

In *Drosophila*, the EB is generated by R neurons. Their layer-specific projections form a toroidal neuropil located dorso-anteriorly straddling the midline of the protocerebrum^29–31^. Based on their layer-specific projections, R neurons are classified into four subtypes (R1-4) each resolved by subtype-specific Gal4 lines^30,31^ (Fig. 1 and Supplementary Fig. 1). In order to reconstruct the origin, formation and circuit assembly of the EB, we first examined *pox neuro (poxn)*, which is expressed in adult R neurons^14^ as well as in embryonic basal forebrain lineages that co-express *Engrailed*^32^ (*en*) (Supplementary Fig. 2).

Protocerebral *poxn*+ lineages derive from embryonic stem cells, neuroblasts ppd5 and ppd8, that are distinguished by *en* and *Dachshund* (*Dac*) expression^33^, with *Dac* restricted to ppd8 (Fig. 2a-f). To determine whether adult EB *poxn*+ cells share their origin with embryonic lineages, we used genetic tracing^34^ to identify lineage relationships from the time of embryogenesis up to the formation of the adult brain employing a heritable membrane-tethered fluorescent marker (Fig. 2g).

*en-Gal4* tracing labelled R neurons that are immunopositive for anti-Poxn (Fig. 2h). Similarly, *poxn-Gal4* tracing identified *poxn>tub>mCD8::GFP* labelled R neurons and their neuropil, all of which are immunopositive for anti-Poxn (Fig. 2i). In contrast, *Dac>tub>mCD8::GFP* did not label R neurons and showed no overlap between protocerebral *poxn*+ cells and membrane-tethered *GFP* (Fig. 2j). Tracing with *en-Gal4, poxn-Gal4* and *Dac-Gal4* recapitulated endogenous *en*, *poxn* and *Dac* expression. We never observed *de novo* expression coinciding with *poxn* protocerebral cells, nor did we detect any evidence for cell migration into this basal forebrain region. Additionally, GFP labelling of subtype-specific Gal4 lines together with genetic tracing and anti-Poxn immunolabeling identified *poxn* expression in GFP-labelled R neuron subtypes (Fig. 2k-m and Supplementary Fig. 3a-d). In contrast, we did not detect anti-Dac immunoreactivity in *poxn*+ cells (Supplementary Fig. 3e-n). Because Dac expression distinguishes between neuroblasts ppd5 and ppd8, our data identify R1-4 neurons as lineage-related sister cells derived from a pair of bilateral symmetric stem cells, embryonic neuroblast ppd5.

**Figure 2.**
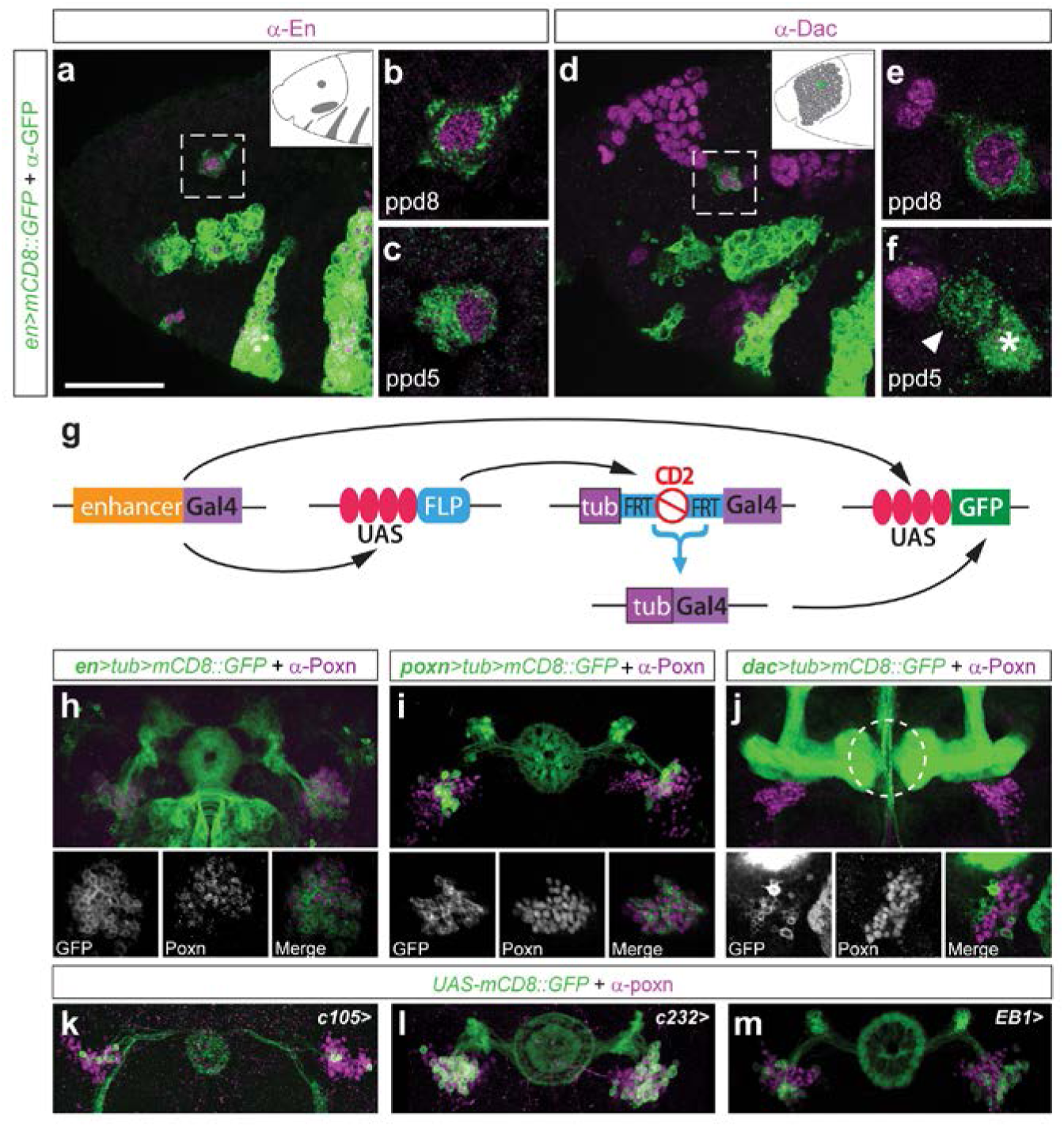
Ellipsoid body R1-4-neurons derive from embryonic neuroblast ppd5. (**a-f**) Confocal images of stage 11 embryos, (anterior, left; dorsal, up; see inset cartoons in a,d; dashed box in a enlarged in b,c; dashed box in d enlarged in e,f). *en>mCD8::GFP* visualises engrailed expression domains overlapping with endogenous *en* in procephalic neuroectoderm, including neuroblast ppd8 (b,e) and ppd5 (c,f). Ppd5 and ppd8 are distinguishable by Dachshund (Dac) expression restricted to ppd8 (d,e). (**g**) Genetic tracing system; Gal4 activates UAS-GfP and UAS-FLP which in turn excises stop cassette thereby activating constitutive active tub-Gal4 labelling all progeny derived from founder cells where excision event took place. (**h-j**) Lineage tracing of *en, poxn* and *Dac* expressing central brain lineages. Confocal images of *en>tub>mCD8::GFP* (**h**) and *poxn>tub>mCD8::GFP* (**i**) show derived lineages including tangential ellipsoid body R neurons and their neuropil; all GFP+ cells express Poxn (small insets). (**j**) *Dac>tub>mCD8::GFP* labelling excludes R neurons; GFP+ cells do not express *poxn* (insets). (**k-m**) R1-specific *c105-Gal4>*, R3-specific *c232-Gal4>* and R2/4-specific *EB1-Gal4>UAS-mCD8::GFP* (green) visualizes cells and subtype-specific, layered projections in EB neuropil. Anti-Poxn immunolabelling (magenta) reveals *poxn* expression in all R neuron subtypes R1-4. Scale bars: 50μm.

### R neurons form a connectivity network among sister cells

EB cells multiply during development and, similar to observations on grasshopper CX development^35^, establish an orthogonal axon scaffold that is later reshaped by morphogenesis into the EB’s characteristically toroidal architecture^29–31^. To determine R neuron connectivity, we utilized the protocerebral *poxn* enhancer element^14^ and generated *poxn-Gal4* strains that either target subsets or the majority of *poxn*+ R neurons (Supplementary Fig. 4a,b). This allowed us to determine their connectivity pattern. We combined *poxn-Gal4* (Fig. 3), or R1-4-specific Gal4 lines *c105, c232* and EB1(Fig. 4a-d) with the presynaptic marker *synaptotagmin::GFP* and the postsynaptic marker *DenMark^16^.* Co-labelling revealed apposed *Syt::GFP* and *DenMark* punctae across layers and modules (Fig. 3b-d and Fig. 4a-d; Supplementary Movie 1). Super-resolution microscopy with structured illumination confirmed *Syt::GFP* and *DenMark* labelled apposing punctae in EB R neuropil (Supplementary Fig. 4c and Supplementary Movie 2). Clonal analysis identified ramified terminal synapses densely distributed along R neuron subtype- and layer-specific arborisations (Fig. 4e). These data indicate that *Syt::GFP, DenMark* labelled apposing punctae visualize synaptic connections of lineage-related R neurons (Fig. 4f).

**Figure 3.**
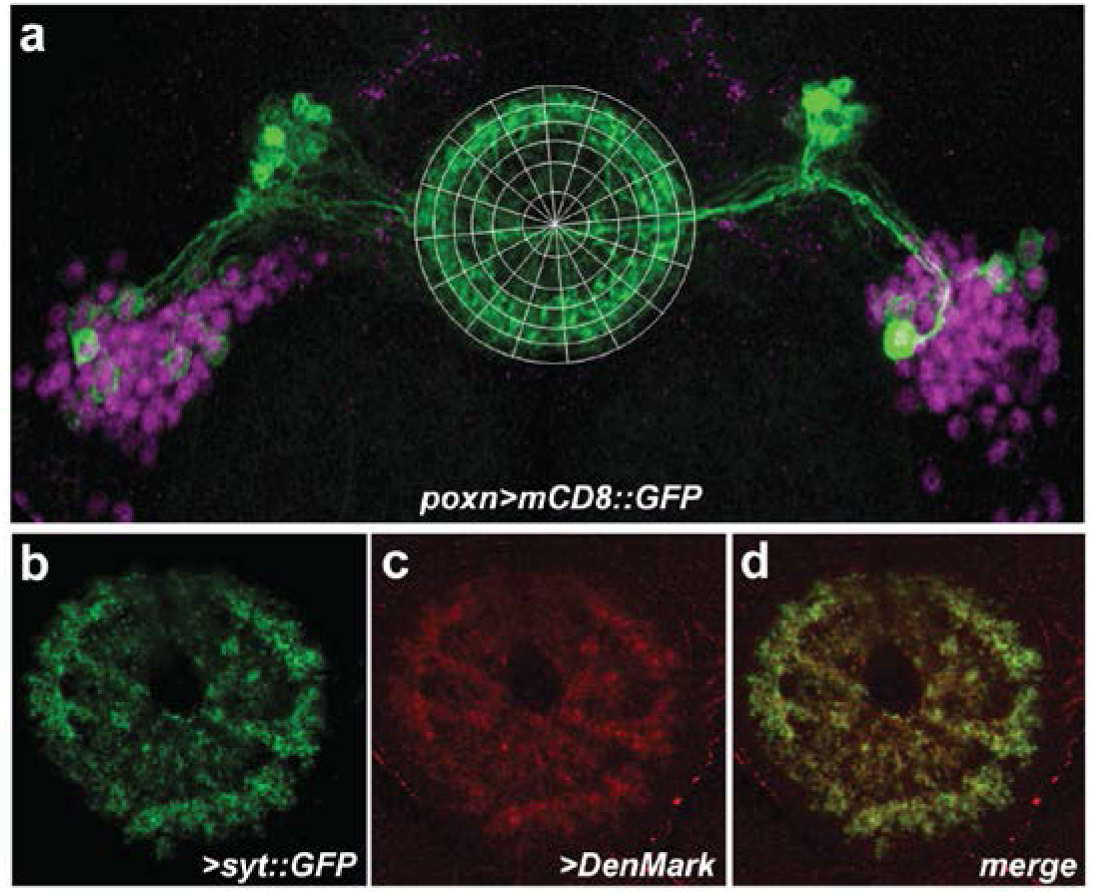
Synaptic targeting of lineage related R neurons. (**a-d**) Confocal images of *poxn^brain^-Gal4* expressing *UAS-mCD8::GFP* in subsets of tangential R neurons. (a) *poxn^brain^>mCD8::GFP* immunolabelled with anti-poxn reveals that GFP positive cells express endogenous Poxn. (b) Presynaptic *poxn^bram^>Syt::GFP* visualises EB neuropil. (c) Postsynaptic *poxn^brain^>DenMark* also visualises EB neuropil. (d) *poxn^brain^>Syt::GFP, DenMark* identifies GFP/DenMark-labelled, apposed punctae in the EB neuropil.

**Figure 4.**
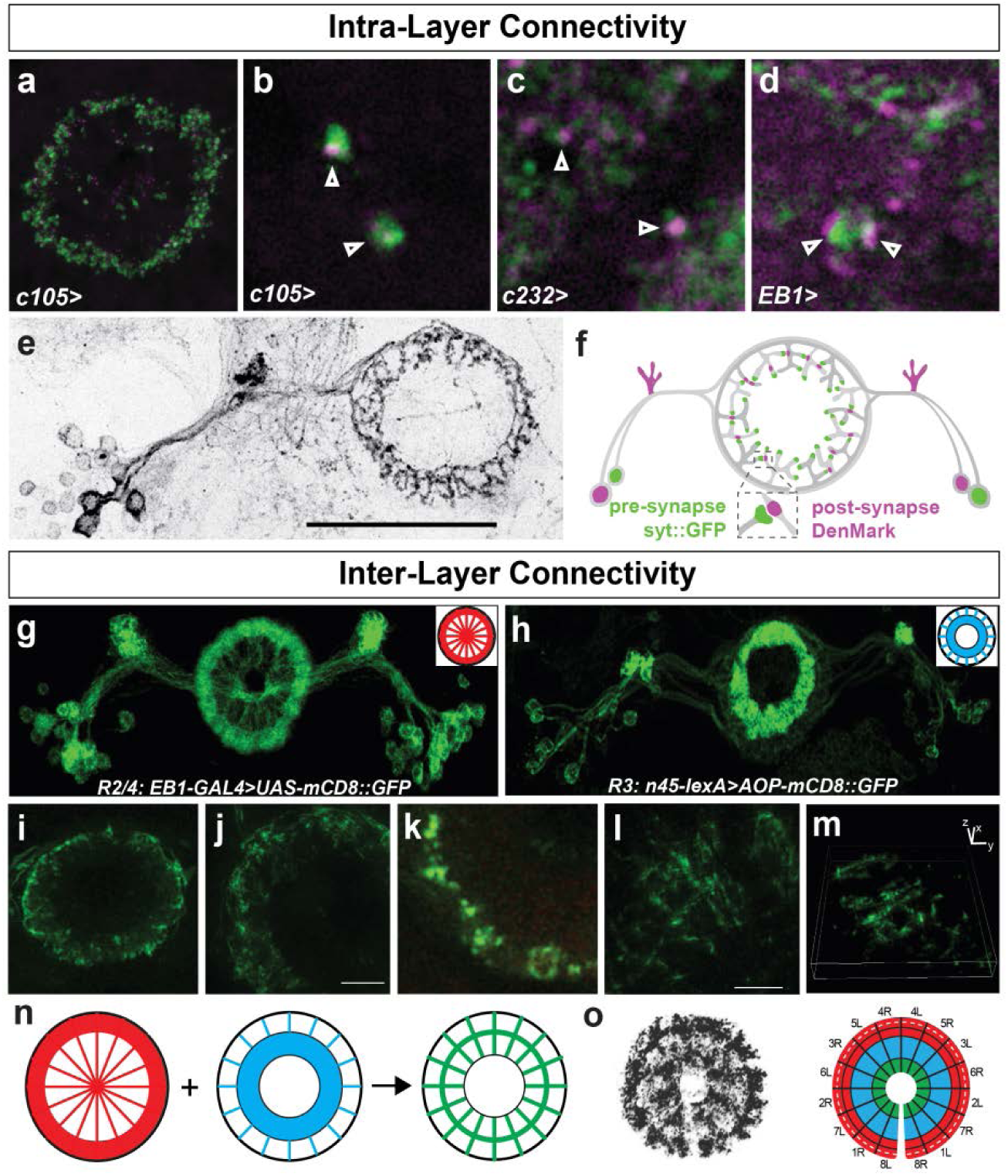
EB R neurons synapse onto lineage-related sister cells. (**a-e**) Genetic labelling of pre- and post-synaptic compartments of EB R neurons. (a-d) Confocal microscopy of R1-4 neuron-specific Gal4 driven presynaptic *UAS-syt::GFP* (green) and postsynaptic *UAS-DenMark* (magenta) visualising apposed punctae in EB neuropil (arrowheads). (**e**) Clonal labelling of R2/4 neuron reveals layer-specific synapses spanning entire circumference of EB neuropil. (**f**) Cartoon of *R Gal4>Syt::GFP, DenMark* labelled apposed punctae indicating R neuron synapses. (**g-m**) *GFP reconstitution across synaptic partners (GRASP)* using (g) *EB1-Gal4>mCD8::GFP* (inset, *EB1-Gal4* pattern in ring neuropil), and (h) *NV45-LexA::VP16>LexAop-mCD8::GFP* (inset, *NV45-LexA* pattern in ring neuropil), (***i-m**) Multi-photon z-stacks of NV45-LexA:: VP16/LexAop-CD4::spGFP11;EB1-Gal4>UAS-CD4::spGFP1-10 identifies reconstituted GFP in punctate and elongated, grid-type pattern. (**m**) 3D reconstruction of several confocal stacks of NV45-LexA::VP16/LexAop-CD4::spGFP11;EB1-Gal4>UAS-CD4::spGFP1-10, (n) Cartoon illustrating EB ring projections of EB1- Gal4, NV45-LexA and their combination with GRASP, colour coded as in Fig, 1c, (**o**) poxn^bram^- Gal4* driven synaptic targeting visualises terminals in all layers and modules of EB ring neuropil (left panel, see also Fig. 3a), thus covering all segments of sensory space represented in the EB (right panel, see also Fig, 1c), Scale bars: 5μm.

To gain further insights into EB connectivity we next performed electron microscopy analyses of the EB neuropil. At low-resolution (Supplementary Fig. 4d) dense neuronal profiles are resolved in any segment of the EB, each encompassing an outer and two inner domains, which define the concentric arrangement of modules (also called ‘wedges’^26^). Supplementary Figure 4e demonstrates the location of synapses (right panel, in purple), which across sections allows an estimate of synaptic densities. Our analysis revealed that domains of the inner EB ring have smaller profiles with several presynaptic sites, many of which appose large profiles, also possessing presynaptic sites (Supplementary Fig. 4e,f). In addition, we also detected pairs of synapses opposing larger boutons, thus revealing synaptic convergences (Supplementary Fig. 4g). More rarely, tetrad synapses terminating onto four postsynaptic profiles contain dense core vesicles, indicating that neuromodulatory elements likely participate in EB functionality (Supplementary Fig. 4h).

To further substantiate connectivity among R neurons, we used *GFP reconstitution across synaptic partners (GRASP).* GRASP utilizes the expression of two split-GFP halves that only fluoresce when reconstituted^37^. To visualise potential connections between R neurons, we used split-GFP fragments *UAS-CD4::spGFP1-10* and *LexAop-CD4::spGFP11 (ref. 38)*, combined with R2/4-specific *EB1-Gal4* (Fig. 4g and Supplementary Fig. 5a) and *NV45-LexA::VP16* specific to R3 neurons (Fig. 4h and Supplementary Fig. 5b). When *EB1-Gal4* and *NV45-LexA* were combined for GRASP, reconstituted GFP fluorescence was detectable around the EB’s circumference and intersecting modules, with most intense GFP signal observed at the intersection of R3 and R2/4 layers (Fig. 4n). However reconstituted GFP fluorescence was not detected in respective controls (Supplementary Fig. 5d). In contrast, super-resolution microscopy of GRASP signals revealed punctate (Fig. 4i-k) and elongated, grid-type GFP expression extending across R2-R4 layers and modules (Fig. 4l,m and Supplementary Movies 3 and 4). Of note, high-resolution confocal microscopy of control *EB1-Gal4>UAS-mCD8::GFP* revealed not only intense labelling of R2/4 layers but also spoke-like projections towards the inner ring (Fig. 4g inset and Fig. 4n), while *NV45-LexA>AOP-mCD8::GFP* controls revealed intense labelling of R3 layers but also projections within the outer ring (Fig. 4h inset and Fig. 4n). When compared to control patterns (Fig. 4g,h), the detected GRASP signal thus also suggests inter-layer connectivity between EB1 and NV45 (Fig. 4l-n). Importantly, the suggested connections were observed with confocal and super-resolution microscopy not only for specific layers, as visualised for example with *c105>Syt::GFP, DenMark* (Fig. 3a,e) and *EB1>Syt::GFP, DenMark* (Supplementary Fig. 4c), but also for modules as seen with *poxn>Syt::GFP, DenMark* (Fig. 3a-d and Fig. 4o). Taken together, these data suggest connectivity among R neurons that extends across layers and modules of *the EB neuropil*.

### GABAergic inhibition of R neurons

We next investigated physiological properties of the R neuron circuitry. Previous studies indicate a majority of R neurons are GABAergic^29,39^. We therefore tested whether *poxn*+ R neurons are GABAergic. GABA expression was identified in 76% of *c105* R1 neurons, 70% of *c507* and 93% of *c232* R3/4d neurons, 73% of *c819* and 94% of *EB1* R2/4m neurons (Supplementary Table 1). These data suggest that *c232* and *EB1* R neurons are inhibitory GABAergic interneurons. To test this hypothesis, we carried out functional imaging using the bioluminescent photoprotein GFP-aequorin (GA)^40^. We expressed GA in c232 neurons and measured photon emission related to calcium-induced bioluminescence activity in open-brain preparations (Fig. 5a,b). Direct bath application of 250□3n picrotoxin, a noncompetitive blocker of GABA_A_ receptor chloride channels, resulted in photon emission (up to 300 photons/s) that persisted for several minutes (Fig. 5c). These data suggest that picrotoxin caused a widespread disinhibition of GABAergic c232 neurons.

**Figure 5.**
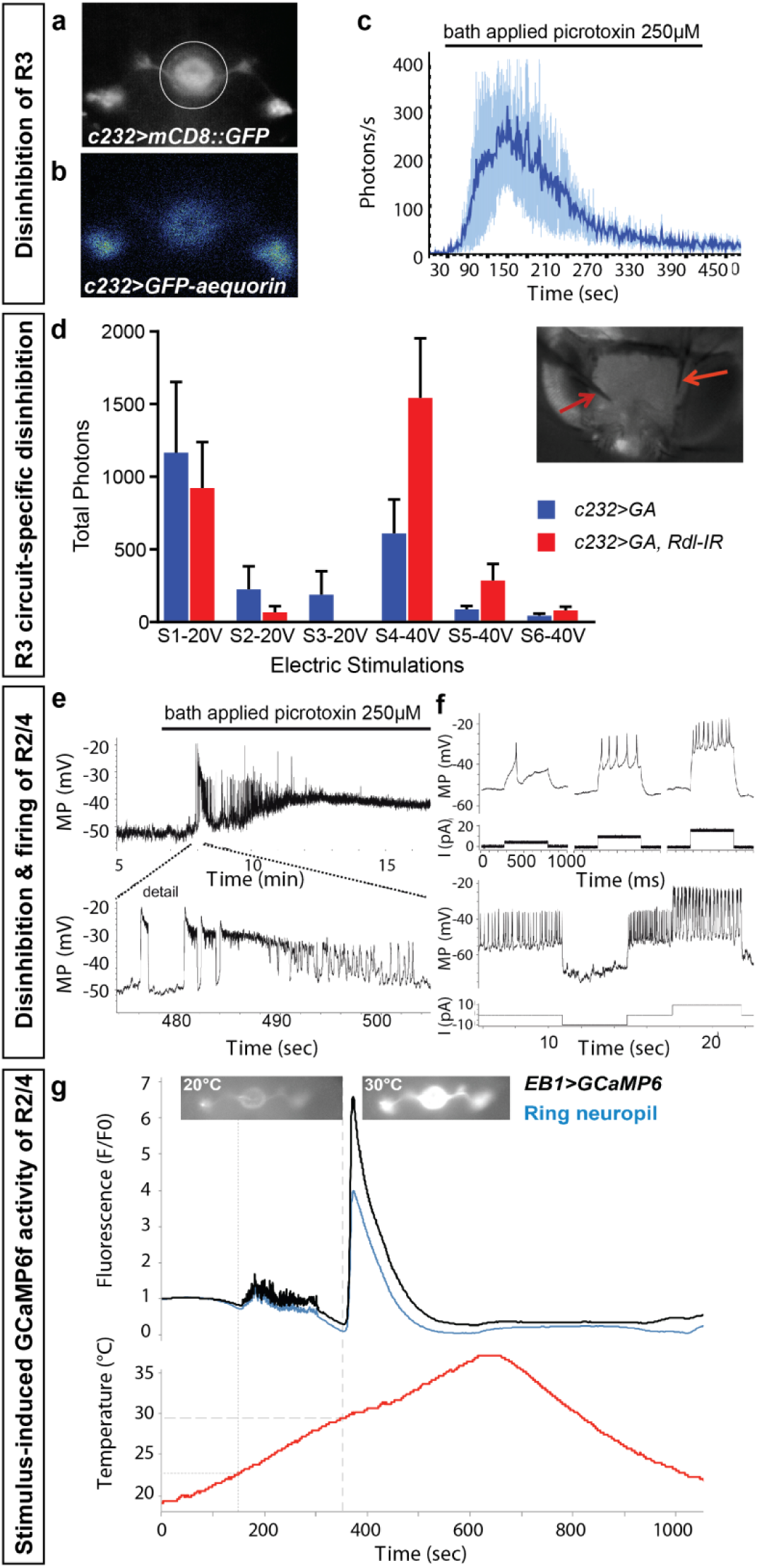
Physiological properties of inhibitory R2-4 neurons. (**a-d**) Calcium-dependent functional imaging of R neuron activity using GFP-aequorin (GA). (**a,b**) *c232>GA* showing expression of bioluminescent photoprotein GA in c232 R neurons (a, GFP; b, photon signal). (**c**) Photon emission in *c232>GA* after picrotoxin application, indicating inhibitory GABAergic activity of R3 neurons. (**d**) c232 circuit- and layer-specific knock down of *Receptor of dieldrin (Rdl)* combined with expression of *UAS-GA* (*c232>GA, Rdl-IR*). Activity measurements in open brain preparations (inset, electrodes highlighted with red arrows) after 20V and 40V electric stimulation, each separated by 2min. Note significant circuit and layer-specific increase of photon emission in *c232>GA*, *Rdl-RNAi* with stimulation protocol 1 at V40, indicating that GABAergic c232-neurons are inhibited by c232-mediated GABA-A receptor signalling. (**e, f**) Electrophysiological recordings of R neurons. Whole-cell recordings of *EB1>mCD8::GFP* R2/4-neurons (n=20). (**e**) Spiking and bursting after picrotoxin application (n=6). (**f**) Stimulus-response of EB1 neurons caused by current injection. MP, membrane potential; I, current. (**g**) GCaMP6f imaging of EB1-Gal4 specific R2/4-neurons during increasing temperature ramps (n=9). Note that sparse activity starting by 23°C (tight dashed line) resolves in GCaMP6f spike around 30°C (wide dashed line) for both cell bodies of R2/4-neurons (black line) as well as their ring neuropil (blue line). Mean ± SD.

To corroborate these observations, we carried out EB circuit- and layer-specific knock down of the *Drosophila* GABA_A_ receptor, *Receptor of dieldrin* (*Rdl*) and combined it with expression of *UAS-GA (c232>GA, Rdl-IR).* To simulate input activity of varying strength, we implanted electrodes into open-brain preparations (Fig. 5d, inset) and injected current of 20V in three stimulations, followed by another three 40V stimulations, each of which lasted 5ms separated by 2min rest (Fig. 5d). When compared to *c232>GA* controls, 40V stimuli but not 20V stimuli resulted in increased photon emission in *c232>GA, Rdl-IR* (Fig. 5d; p<0.01 two-way ANOVA, n=14). These data suggest that the circuit-specific Rdl knockdown increased the electric stimulus induced-response of c232 neurons when compared to *c232>GA* controls. Moreover, consistent with a disinhibitory effect of *c232>Rdl-IR*, the observed GA activity recordings suggest c232 circuit-specific inhibition among c232 neurons.

To gain further insight into physiological properties of GABAergic R neurons, we performed whole-cell patch recordings from GFP-labelled *EB1* R neurons. These revealed resting membrane potentials of −52.9 ± 4.0mV (mean ± standard deviation, SD) and input resistances of 1185 ± 407MΩ (n=20). Following bath-application of 250μM picrotoxin, all of the six recorded neurons exhibited individual spikes followed by bursting activity that lasted between 10-40s (Fig. 5e). Bursts (0.5-10Hz frequency) comprised up to 60 individual spikes, with an instantaneous spike frequency ranging from 30-100Hz, followed by strong depolarisation which suppressed further activity. To test their stimulus-salience response, we injected increasing amounts of depolarising current (up to 40pA) into *EB1* R neurons, which evoked repetitive firing with successively more spikes triggered by higher currents, up to a maximum frequency of 80Hz (Fig. 5f), suggesting that spike frequency increased proportional to stimulus salience/strength. Together, these data suggest that *c232* and *EB1* R neurons are inhibitory GABAergic interneurons that are inhibited by GABA_A_ receptor signalling.

### Sensory-motor transformation by inhibitory R neurons

Ellipsoid body R neurons have been shown to play a role in goal-directed locomotion, spatial orientation learning and memory, as well as in sleep, attention, arousal, and decision-making^13,14,16–18,20–26^. All of these observations suggest that these neurons are involved in the assessment of sensory information leading to the determination of a behavioural action. To further address this, we made use of temperature-related behaviour which in *Drosophila* is known to elicit goal-directed locomotion^41^ (Fig. 6).

**Figure 6.**
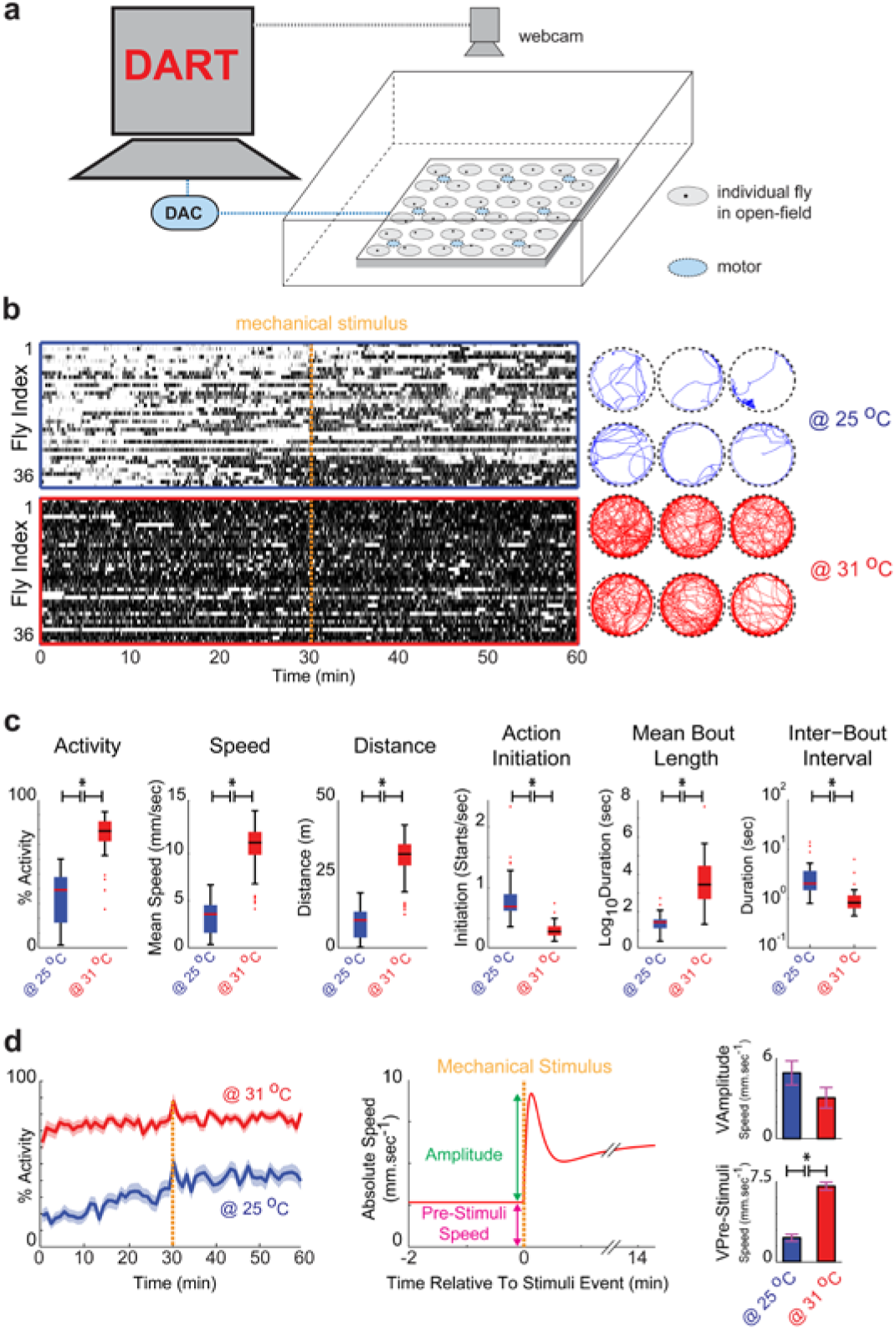
Open-field assay for quantitative analysis of motor activity and arousal state. (**a**) Behavioural setup with arenas in temperature-controlled incubator. Modified DART software^70^ was utilized for video-assisted motion tracking which also controls mechanical stimulation (vibration) via a digital to analogue converter (DAC). Multiple motors represented by the light blue ovals are connected to the DAC via a simple circuit board. (**b**) Left, raster plots showing sequences of activity (black bar) and inactivity (white space between black bars) of 36 control flies *w^1118^* recorded either at 25°C (top panel surrounded by blue line) or 31°C (bottom panel surrounded by red line); vertical dashed orange line indicates mechanical stimulation. Right, trajectories shown for 6 flies recorded for the first 10min either at 25°C (in blue) or 31°C (in red). (**c**) Parameters deduced from motion tracking detailing motor actions and their organisation into action sequences; mean activity, speed and distance travelled while flies are active; action initiation = number of starts; mean bout length = mean duration/maintenance of activity bout; inter-bout interval = mean duration of pauses between bouts of activity. (**d**) Arousal state measured as response to mechanical stimulation; left, average speed over the time-course of the experiment with stimulus represented by dashed orange line; middle, pre- and post-stimulus speed used to derive amplitude of stimulus response as proxy for arousal state; right, quantification of baseline speed (V_Pre-Stimuli_) and the amplitude of the response (V_Amplitude_).

Temperature sensation in *Drosophila* is mediated by thermal receptors, with sensors for innocuous warmth located in the brain^42^. Four warmth-activated thermosensory neurons called Anterior Cells (AC) express Transient Receptor Potential A1 (TRPA1) essential for temperature integration and the selection of preferred temperatures; these neurons send connections to the subesophageal ganglion, to two antennal lobe glomeruli and to the superior lateral protocerebrum (SLP)^43,44^. The SLP is innervated by Fm-neurons of the FB^45^ which itself is connected with the EB (Fig. 1b). Since none of the known thermal receptors are expressed in the EB, we asked whether integrated temperature information is received and processed by R neurons.

To resolve this question, we expressed GCaMP6f (ref. 46) in *EB1* R neurons. GCaMP-fluorescence of preparations exposed to temperatures ramps ranging from 18°C-35°C showed a response in 7 out of 9 brains (Fig. 5g). GCaMP-fluorescence detection revealed a saccade of small peaks starting around 23°C, indicative of activation of individual neurons, which by >30°C focused into fluorescence spiking for both *EB1>GCaMP6* expressing neurons and their ring neuropil (Fig. 5g). Spiking occurred instantaneously and across the whole population, with GCaMP-fluorescence intensity 4-7-fold above baseline for ring neuropil and cell bodies, respectively, and rapidly declined over 2 minutes, followed by inactivity even at noxious 35°C (Fig. 5g). The pattern and time course of this calcium increase resembles the steadily increasing activity seen in electrophysiological recordings in response to application of picrotoxin (Fig. 5e) or to increasing amounts of depolarizing current (Fig. 5f). These results suggest that *EB1* R neurons assess stimulus input of sensory information, such as temperature changes, by threshold-dependent neuronal activity.

### GABAergic inhibition of R neurons mediates motor actions

The selection and gating of behavioural actions comprise their initiation, maintenance and termination as well as their assembly into activity sequences, which together generate adaptive behaviour^9^. The most basic selection is between action and inaction. The failure of such selectivity is illustrated by pathologies of human motor control and motivational disorders, such as the indecisiveness of motor actions seen in Parkinsonism, and the diminution of voluntary motor actions seen in abulia^47^. These behavioural symptoms exemplify disease-specific alterations in the initiation, maintenance or termination of actions and their goal-directed assembly into action sequences. Analogous pathologies occur in *Drosophila.* These are characterized by anatomical alterations or defective neuromodulation of CX circuits^11,18^. Given its proposed role in action selection, EB dysfunctions are likewise expected to affect, for example, the frequency in which flies initiate and maintain behavioural actions, such as bouts of walking.

We measured such behavioural alterations using open-field assays^48^, employing video-assisted motion tracking to record freely moving *Drosophila* (Fig. 6a). Flies were kept in 35mm diameter arenas and after 1hr of adaptation were recorded for 60 minutes, with a short pulse of mechanical stimulation applied after 30 minutes (see supplementary methods). Based on these recordings, we generated raster plots and trajectories for each individual fly and extracted parameters detailing their behaviour including action initiation and maintenance and their response to sensory stimulation (Fig. 6b-d), each of which are integral to the action selection process49. To examine R neuron functions, we utilized *c105, c232* and *EB1-Gal4* drivers to express specific UAS-transgenes targeted to R1, R3 and R2/4 neurons, respectively (Supplementary Fig. 1). We first expressed *UAS-Rdl-IR* to knock down the *Drosophila* GABA_A_ receptor, *Rdl*, which identified alterations for R neuron subtypes (Fig. 7 and Supplementary Table 2). *Rdl-IR* expression resulted in increased activity and walking distance, which for R1 and R3 was accompanied with less frequent initiation of actions but their increased bout length. *EB1>Rdl-IR* flies were also faster and paused less. Mechanical stimulations did not reveal impaired behavioural responses but an increase in the amplitude was observed for R1 and R3 that could be attributed to decreased pre-stimuli speed (Supplementary Fig. 6). These data suggest that GABAergic inhibition of R neurons coordinates voluntary motor actions and their assembly into action sequences.

**Fig. 7.**
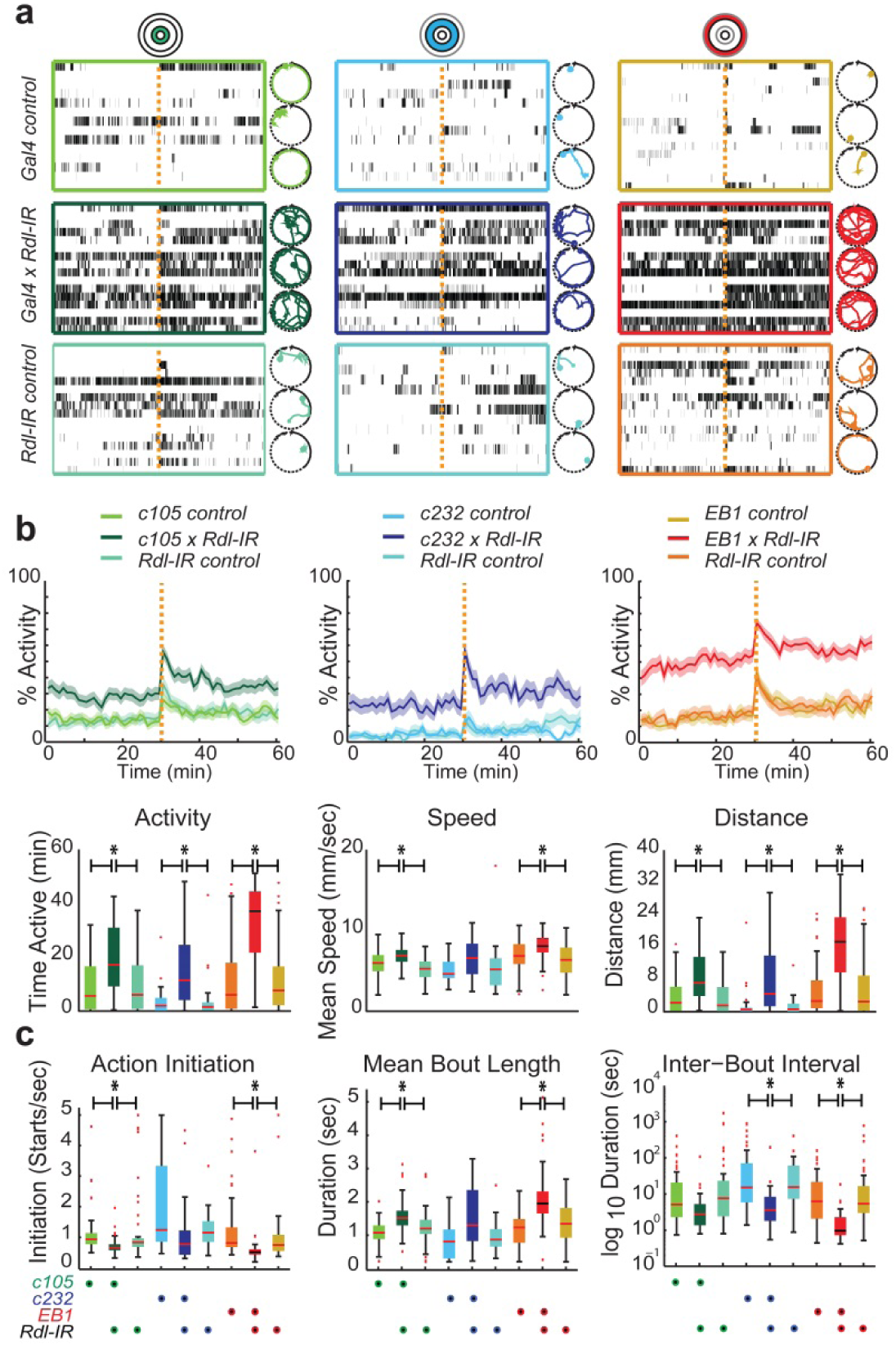
Inhibitory GABAergic activity of EB R neurons in motor action selection. Behavioural measures of R1, R3 and R2/4 neuron subtype-specific expression of *Rdl-IR* targeting the *Drosophila* GABA_A_ receptor, *Receptor of dieldrin* (*Rdl*), and respective controls recorded at 25°C. (**a**) Raster plots and 10min movement trajectories are shown for respective EB ring layers (colour coded). (**b**) Top row, average activity over 60min recordings, including response to mechanical stimulation given after 30min (orange dashed line); bottom row, action measures of mean activity, speed while active and distance travelled. (c) Action sequence measures of mean number of starts per sec (action initiation), mean duration of an action (mean bout length), and mean duration of pauses in between actions (inter-bout interval). Mean ± Standard Error of the Mean (SEM), asterisks indicate p<0.05.

### R neurons mediate the selection and maintenance of behavioural actions

A Action selection circuitry is readily engaged by external stimuli that drive an animal’s behaviour^49^. Previous studies identified a key role of the insect EB in a variety of behavioural activities relating to sensory stimuli^16–18,20–22,26,27^, suggesting that stimulus-induced action selection requires EB function. We hypothesized that modifying EB R neuron circuitry should result in an acute impact on behaviours that relate to salient external stimuli. To test this we focused on temperature-related behaviour in the knowledge that *Drosophila* actively avoid temperatures above 29°C, an aversive temperature that readily elicits goal-directed locomotion^41^ (Fig. 6b and Supplementary Fig. 7a). To utilise this activity in our analysis of the role of R neurons, we expressed either *Tetanus-Toxin-Light-Chain (TNT)*, encoding an inhibitor of synaptic transmission^50^, or *NaChBac* encoding a bacterial depolarisation-activated sodium channel^51^ to increase neuronal excitability (Supplementary Fig. 8). In order to rule out any potential developmental effects, we also utilized thermogenetic activation using the cation channel *TRPA1* (ref. 43) to activate neurons or the dynamin mutant *Shibire^TS^* to block vesicle endocytosis^52^.

We first determined thermal preference behaviour for *c105>TNT, c232>TNT* and *EB1>TNT* flies, which revealed peaks of around 25°C (Supplementary Fig. 7a) as preferred temperature^41^. In addition, we ruled out that R neurons regulate fixed action patterns such as startle-induced negative geotaxis (SING) which triggers an innate escape response^53^, or that *R neuron>TNT* flies were impaired in leg coordination, or subject to motor abnormalities, or to muscle fatigue. Except for a single meta-thoracic neuron cluster in *c105*, none of the lines show Gal4 expression in muscles or motor neurons (Supplementary Fig. 1), and SING analysis revealed no significant differences (Supplementary Fig. 7b), which would be the case if motor neurons or leg coordination were affected^53,54^.

Next we analysed open-field behaviour of *c105, c232* or *EB1-Gal4* flies misexpressing *UAS-TNT* or *UAS-NaChBac*. Recordings at 31°C revealed behavioural alterations of *c105>TNT* flies that initiated more, albeit slower, actions of shorter duration (bout length) which were intermitted by longer pauses (Interbout Interval, IBI length); these alterations, however, were unrelated to their arousal state which remained unchanged (Fig. 8 and Supplementary Fig. 9). In contrast to R1-related behavioural alterations, we did not observe any significant changes relating to R2-R4 neurons. This R1 subtype-specific modulation of behaviour was largely recapitulated by misexpression of *Shibire^TS^* (Supplementary Fig. 10). No significant alterations in the behaviour of *>NaChBac* expressing flies were observed at 31°C (Supplementary Fig. 11). In contrast, TRPA1-mediated acute activation of EB-R neurons at 31°C identified behavioural alterations almost exclusively restricted to R3 neurons (Fig. 9). Thus, *c232>TRPA1* flies were characterized by significantly decreased basal activity that contrasted with more actions initiated which were not maintained very long and intermitted by extended pauses; these behavioural alterations were unrelated to changes in the arousal state of *c232>TRPA1* flies, which remained unaltered (Supplementary Fig. 6). Together these data establish specific roles of R neuron activity in the selection and maintenance, and thus gating of behavioural actions.

**Figure 8.**
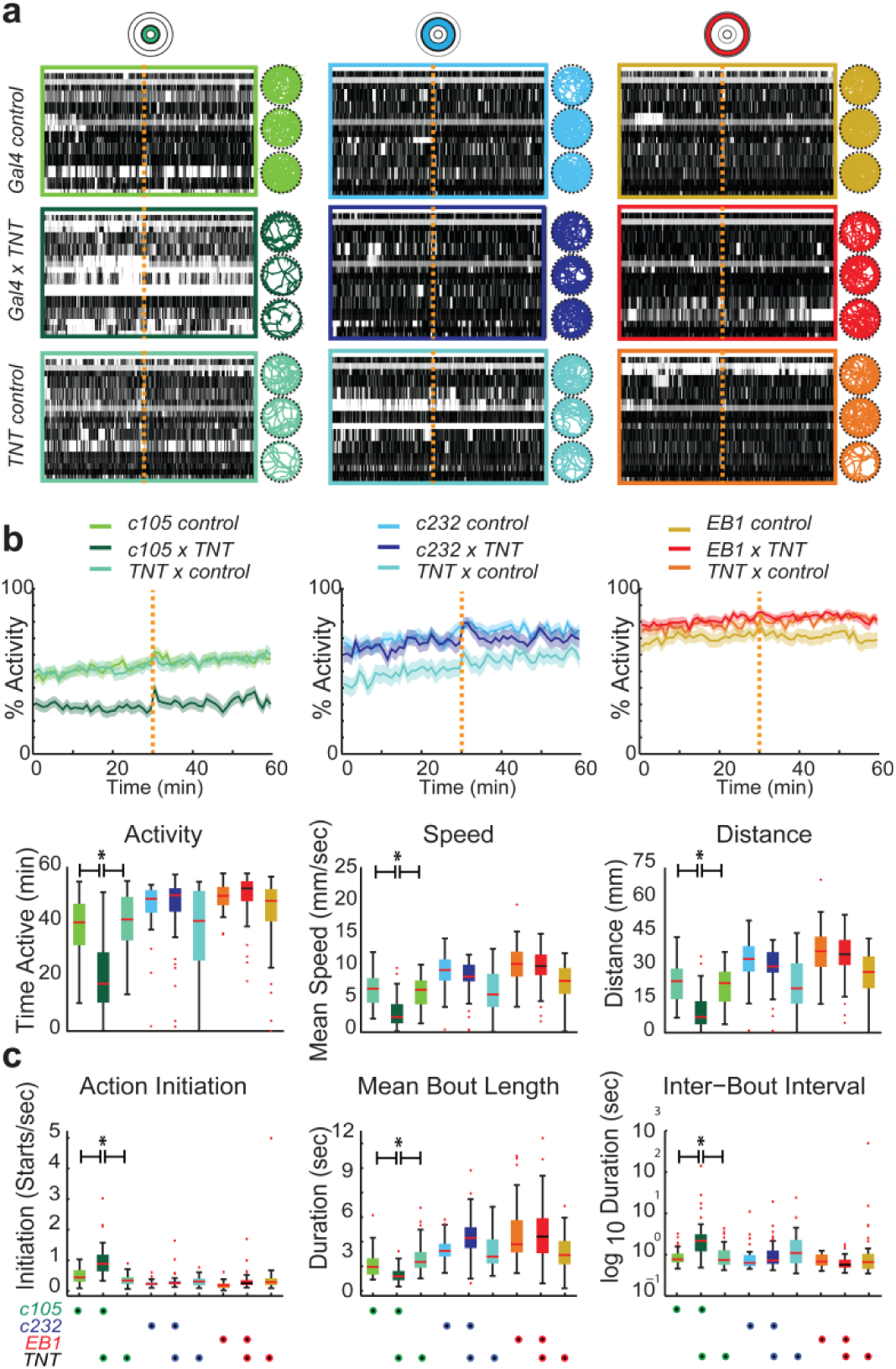
R1-neurons mediate motor output of the EB microcircuit. Behavioural measures of R1, R3 and R2/4 neuron subtype-specific expression of *Tetanus-Toxin-Light-Chain* (*TNT*), encoding an inhibitor of synaptic transmission^50^, and respective controls recorded at 25°C. (**a**) Raster plots and 10min movement trajectories are shown for respective EB ring layers (colour coded). (**b**) Top row, average activity over 60min recordings, including response to mechanical stimulation given after 30min (orange dashed line); bottom row, action measures of mean activity, speed while active and distance travelled. (c) Action seauence measures of mean number of starts per sec (action initiation), mean duration of an action (mean bout length), and mean duration of pauses in between actions (inter-bout interval). Mean ± SEM, asterisks indicate p<0.05.

**Figure 9.**
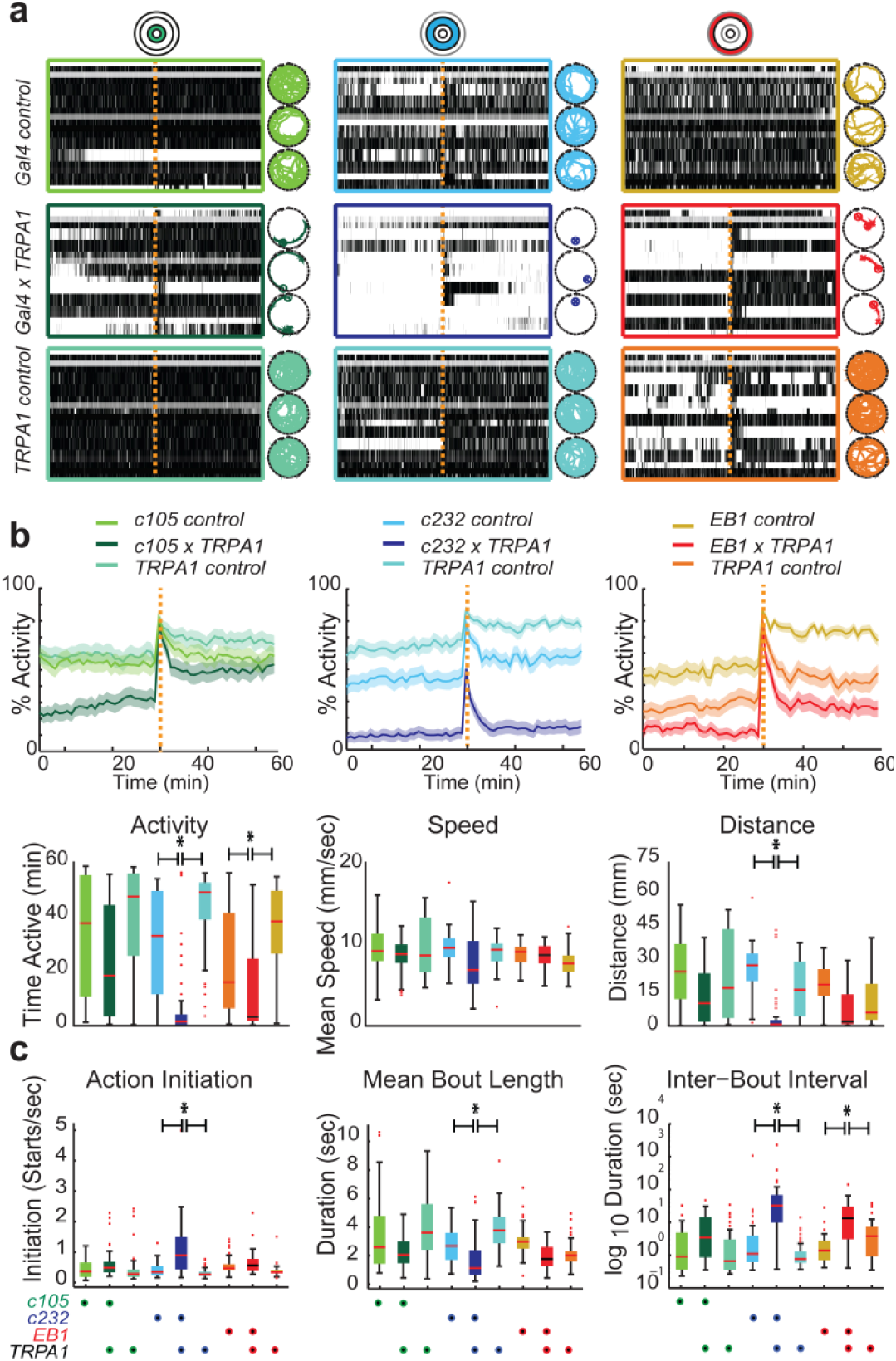
EB R3-neurons mediate motor action selection. Behavioural measures of R1, R3 and R2/4 neuron subtype-specific expression of the cation channel *TRPA1* (ref. 43), and respective controls recorded at 25°C. (**a**) Raster plots and 10min movement trajectories are shown for respective EB ring layers (colour coded). (**b**) Top row, average activity over 60min recordings, including response to mechanical stimulation given after 30min (orange dashed line); bottom row, action measures of mean activity, speed while active and distance travelled. (c) Action sequence measures of mean number of starts per sec (action initiation), mean duration of an action (mean bout length), and mean duration of pauses in between actions (inter-bout interval). Mean ± SEM, asterisks indicate p<0.05.

### A computational model of EB action selection circuitry

The sum of our data supports the proposition that R neurons form GABAergic inhibition circuitry among inhibitory sister cells that underlies the selection and maintenance of behavioural actions. To further explore the computational features of such circuitry within the CX, we developed a neural model relying on leaky integrator units to simulate the mean field activity of neural populations^55^ (see supplementary methods). We constrained the model architecture to replicate known connectivity of the CX, focussing on afferent and efferent EB projections (Fig. 10a). The EB receives and integrates heterogeneous sensory information^10^. For instance, stimulus-induced visual activity in the PB’s and the FB’s columns convey information about a visual feature’s locality in relation to the head and compound eyes^22,26^. In four simulations used as case studies (Fig. 10), noisy information about sensory cues is converted into the activation of PB modules. The features characterizing these stimuli, such as signal intensity and localization in space, are then propagated in parallel to EB modules, activating them dependent on the strength of the signal, as has been shown for visual cues^26^, and weighed by the relative connections. Our present study suggests GABAergic R3 neuron projections are connected to GABAergic R2/4 neurons. Therefore we simulated three EB layers (R1, R2/4 and R3), each divided into 8 modules per hemisphere (as for PB, FB and LAL)^45,56–58^ that are characterized by undifferentiated lateral inhibitions across layers and within each layer (Fig. 10a).

**Figure 10.**
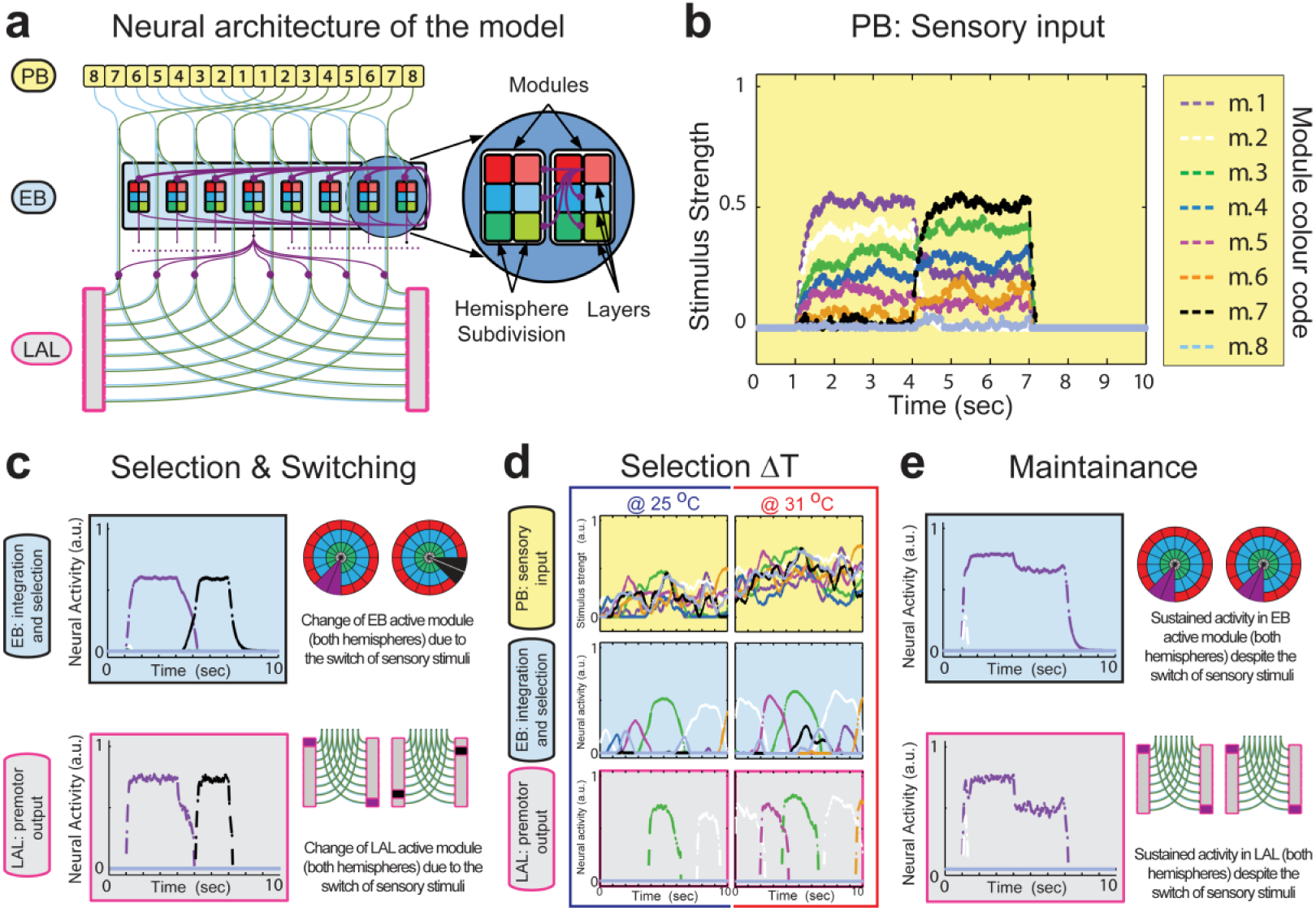
Computational model of the R neuron action selection circuitry. (**a**) Schematic architecture showing organization and connectivity among eight modules per hemisphere of protocerebral bridge (PB), ellipsoid body (EB) and lateral accessory lobes (LAL); note FB is omitted for clarity. Excitatory connections are shown in blue/green (starting from PB); inhibitory connections in magenta (starting from EB). EB colour code (red-blue-green) illustrates concentric layers formed by R1, R3 and R2/4 neuron projections; inhibitory connections among layers and modules highlighted in enlarged circle. (**b-e**) Simulated activity is represented by different colours, shown as signal strength propagated along columnar projections, e.g. module in PB propagates signal towards modules in EB and LAL (magenta). (**b**) Sensory input processed in PB consists of two configurations, each lasting 3 seconds. These signals reach modules of both EB and LAL. (**c**) EB integrates and processes incoming signals (top) via lateral inhibitions, suppressing all competitors, cancelling noise, thus detecting and selecting the strongest stimulus (winner-takes-all dynamics). EB activity is represented both as signal strength (single hemisphere, top-left) and in toroidal representation of modules (double hemisphere, top-right). The LAL receives excitatory inputs from PB, with the activity of all modules inhibited by EB activity except for the winning one. LAL module activity is represented as signal strength (single hemisphere, bottom-left) and in a schematic representing both hemispheres of LAL (double hemisphere, bottom-right). Simulated activity shows the system adapting to changes of sensory stimuli, always selecting the strongest input (first magenta and then black), which also enables switching among new salient sensory input. (**d**) Unstable sensory inputs simulating either 25°C (low value random walk, left) or 31°C (high value random walk, right). Simulated activity in the LAL (graphs on the right) shows motor selections, resulting from a continuous competition among modules in the EB (graph in the centre), which significantly increase in the simulated 31°C condition. (**e**) Selection is maintained due to strengthened connections among inhibitory R neurons, even when the set of stimuli changes (magenta).

We tested a range of parameters to explore the robustness of emerging functions (Supplementary Fig. 12). In particular, we focused on whether the known neurophysiology of the EB is consistent with the proposed functional role of a sensory-driven selector. We opted for fixed lateral inhibition between R neurons, while exploring the consequences of several different configurations of excitatory inputs from the PB and FB to the EB, and of inhibitory outputs from EB to LAL (Supplementary Fig. 12). Our simulations show that parallel input connectivity, coupled with internal lateral inhibitions are sufficient to realize sensory integration, to detect the highest saliency, to cancel competing signals and noise, and to adapt to changes in the environment such as novel sensory input (Fig. 10b-e and Supplementary Fig. 12). The first two case studies show dynamics of several noisy competing stimuli encoded in the activation of PB modules (Fig. 10b). These stimuli are propagated to EB modules where lateral inhibitions mediate competition among all active modules. The resulting computation among R neurons mediates salience detection by winner-takes-all functionality and the selection of only one active module, cancelling both noise and competing signals^59^ and yet allowing adaptation to changes in the topography and configuration of new salient sensory cues (Fig. 10c and Supplementary Fig. 12). Our model also predicts that during persistent stimulus input, such as elevated temperature, the EB circuitry enables continued salience detection and the competitive selection of one active module (Fig. 10d). At the end of a winner-takes-all competition, and depending on the synaptic strength of R neuron connections, our simulations show that lateral inhibition can maintain a selected activity even after input changes (Fig. 10e).

## DISCUSSION

Our results identify a pair of bilateral symmetric, engrailed-expressing embryonic neuroblasts, ppd5, that generate all four subtypes of ellipsoid body R neurons (R1-R4) in the central complex of the adult brain in *Drosophila*. Lineage-related R neurons form GABAergic microcircuitry that when dysfunctional impair the selection and maintenance of behavioural actions. The present results identify the organization of EB microcircuitry as a salience detector, using competitive inhibition to amplify different activity levels between modules thereby implementing a winner-takes-all mechanism for selecting action output.

A key feature of action selection is the evaluation and transformation of sensory representations into motor representations^49^. Consistent with previous findings^16–18,20–22,26^, our results demonstrate that R neurons assess sensory information and mediate the coordination of voluntary motor actions and their assembly into action sequences. Our computational model accounts for the observed results of genetic manipulations and the ensuing behavioural observations, suggesting the R neuron circuitry functions in sensory-motor action selection. The predictions of our model are supported by functional imaging studies showing that specific EB module activity correlates with the azimuth of a salient visual cue^26^. The observed activity “bump” was largely confined to one module but its position within the EB ring did not match a retinotopic position of the visual cue. Rather it represented the fly’s orientation relative to the visual landmark, with module-restricted activity switching along the EB toroid according to changes in the fly’s angular orientation aligned to the sensory _cue_^26^.

Layers and modules in the ellipsoid body have been implicated in a variety of behavioural manifestations that identify key roles of the EB in response not only to visual cues for spatial orientation^15,19,22,24,26^ and memory formation^16,17,21^, but also in response to olfactory cues for ethanol-induced locomotion^20^ and tactile cues for arousal^18^ and turning behaviour^23,25^ as well as in relation to sleep drive^27^. Our findings further suggest that R neurons mediate weighing and selection of sensory representations independent of their modality. In support of this, the present functional imaging and electrophysiological data identify threshold-dependent stimulus responses of R3 and R2/4 neurons, the spiking initiation of which was dependent on stimulus salience, and whose spiking frequency was proportional to stimulus strength (Fig. 5). Moreover, the model simulations predict that dependent on synaptic strength, inhibitory R neuron activity can maintain a selected activity even after inputs change. In accordance with this, an EB module-specific activity “bump” has been shown to persist as after a salient visual cue disappeared^26^. Thus, the selection and maintenance of an active module can be regarded as convergence of an internal representation of a behaviourally relevant sensory cue and the agent’s reference to it, such as angular orientation^26^ and movement intent^25,60^. The availability of an EB module-specific activity for a prolonged time can guide successive behaviours and in this sense exhibits essential features of short-term memory^61^, which in the case of visual short-term memory has been shown to require R neuron activity^16^.

The R neuron circuitry is embedded in a connectivity network with the PB, FB and the LAL^45,56–58^, the latter a premotor command centre projecting to descending neurons that innervate central pattern generators executing motor actions^62^. As a node of convergence, situated several synapses downstream of sensory neurons and several synapses upstream of motor ganglia, the position and connectivity of the EB-LAL interface implicates it in the translation of sensory representations into motor representations. Experiments in locusts^63^ and silk moths^64^ identified that LAL modules preserve information about values and saliency associated with selected sensory representations. Our simulations confirm these findings and predict that the EB exerts selection on premotor commands in the LAL by module-specific columnar projections^10,64^ that retain stimulus values by a one-to-one correspondence. As a consequence, inhibitory projections from the EB towards the LAL can reverberate the selection process and perform a surround gating of the LAL. As a result, only one motor command is selected at a time while the release of multiple motor actions are avoided at the same time (Fig. 10c-e and Supplementary Fig. 12).

In line with these predictions, our results show that artificial manipulations of R neurons are themselves sufficient to initiate, increase or abrogate motor actions and their organization into action sequences (Figs 7–9, Supplementary Figs 6 and 9-11). Disinhibited behavioural activity was observed with *Rdl-IR* expression for each of the EB’s concentric layers, which was inversely correlated with action initiation as flies less frequently initiated behavioural activity. A similar inverse correlation was also observed with specific overexpression of TNT, Shibire and TRPA1 in R neurons. Although all EB layers respond to *Rdl-IR*, the resulting behavioural manifestations reveal subtype-specific differences that are also seen for TNT, Shibire and TRPA1. TRPA1-mediated behavioural changes suggest a specific role of R3 and R2/4 neurons in the initiation and termination of motor action, while c105-specific TNT and Shibire expression caused activity changes indicative of an essential role of R1 neurons as the EB output^10^.

A comparable organization into partially segregated functional units also characterizes inhibitory GABAergic striato-pallidal projections that constitute the direct and indirect pathways of the vertebrate BG^10^. In mice, optogenetic manipulations have shown that direct pathway stimulation facilitates motor actions, whereas indirect pathway stimulation decreases motor activity^2^, both of which are cooperatively active during voluntary movement^3,4^. Furthermore, functional imaging of direct and indirect pathways in mice identified their differential yet concomitant activities underlie the parsing and concatenation of action sequences^65–67^. These observations lend support to the prevailing model of BG-mediated action selection; namely, by disinhibition of a selected motor program and inhibition of competing actions^5,7–10,68^.

In *Drosophila*, our genetic manipulations and neural computations identify facilitation, inhibition and disinhibition of motor activity as differential R neuron functions underlying the initiation, maintenance and termination of actions and their organisation into action sequences. These findings suggest that R neuron circuitry exhibits homologies to direct and indirect pathway activities including their disease-related dysfunction^1,10^. Focal lesions and dysfunction of direct and indirect pathways are associated with movement disorders, such as Parkinson’s and Huntington’s disease and dystonia^5,47^, as well as neuropsychiatric disorders including impaired memory formation, attention deficits, affective disorders and sleep disturbances^6–8,69^. Comparable behavioural deficits are seen in flies with altered R neuron function^16–18,20,26^, suggesting homologous pathological manifestations. It stands to reason that evolutionarily conserved mechanisms underlie action selection in health and disease^10^, including the coordination of higher motor control and motivation.

## METHODS SUMMARY

For extended methods see Supplementary Information.

### *Drosophila* stocks and genetics

Flies were maintained on standard cornmeal medium at 25°C in a 12hr:12hr light/dark cycle. For lineage tracing, *w^1118^* (control) and Gal4 strains were crossed to *UAS-mCD8::GFP, tub-FRT-CD2-FRT-Gal4, UAS-FLP/CyO GMR Dfd YFP.* For clonal analysis, lines *y, w, FRT19A;* +; + and *w, FRT19A, tubP-Gal80, hs-FLP; UAS-nucLacZ, UAS-mCD8::GFP / CyO; tubP-Gal4 / TM6B Hu* were used. For GRASP analysis, R2/4-specific *EB1-Gal4* and R3-specific *NV45-lexA::VP16* driver were combined with *UAS-CD4::spGFP1-10* and *LexAop-CD4::spGFP11*, For functional imaging, *UAS-GFP-aequorin* or *20XUAS-IVS-GCaMP6* were combined with R3/4 *c232-Gal4* or R2/4 *EB1-Gal4.* For behavioural experiments, *UAS-Rdl-IR, UAS-TNT, UAS-NaChBac, UAS-Shibire^TS^*, or *UAS-TRPA1* were crossed to R neuron-specific Gal4 lines, and controls generated by crossing driver and responder lines to *w^1118^*. Further details are provided in the extended methods section of *Supplementary Information*.

### Super-resolution and electron microscopy

Structured Illumination Microscope and Multi-photon microscope (Nikon) were used for super resolution microscopy; 3D reconstruction and video assembly was carried out using Nikon software. For electron microscopy, 80 nm sections were cut on a Leica Ultracut S Ultramicrotome and visualized in a Phillips CM12 transmission electron microscope.

### Functional imaging and electrophysiology

Bioluminescence imaging was performed in open brain preparations of tethered flies. GCaMP6 imaging and electrophysiology were carried out on isolated brains maintained in saline solution. Whole cell current clamp recordings were performed using glass electrodes with 8-12 MΩ resistance filled with intracellular solution and an Axon MultiClamp 700B amplifier, digitised with an Axon DigiData 1440A (sampling rate: 20kHz; filter: Bessel 10kHz) and recorded using pClamp 10 (Molecular Devices, CA, USA).

### Behavioural analysis

Single, 3-5 day old flies were placed in an open-field arena, and their behaviour was recorded with a Logitech c920 camera at 10 frames per second. 36 flies, 12 of each genotype, were recorded simultaneously for 1 hour. After 30 minutes, mechanical stimulation was delivered through shaft-less motors (Precision Microdrives) controlled by custom-made DART software^70^. These behavioural experiments were carried out in a temperature-controlled chamber at either 25°C or 31°C. Thermal preference behaviour was carried out using a linear temperature gradient from 18°C to 31°C.

### Neural network modelling

Ellipsoid body R neuron modelling was performed in MATLAB using custom scripts. Network dynamics were captured with a firing rate model based on a leaky integrator defined by a continuous-time differential equation.

## SUPPLEMENTARY INFORMATION

Supplementary Information includes twelve figures, two tables and four movies, as well as Detailed Methods and Supplementary References.

### ACKNOWLEDGEMENTS

This work was supported by the UK Medical Research Council (G070149; MR/L010666/1), the UK Biotechnology and Biological Sciences Research Council (BB/N001230/1), the Royal Society (Hirth2007/R2), Parkinson’s UK (G0714), King’s College Medical Research Trust (JRC 19/2007), the MND Association (Hirth/Mar12/6085; Hirth/Oct07/6233), Alzheimer Research UK (Hirth/ARUK/2012), and the Fondation Thierry Latran (DrosALS) to F.H. N.J.S. received support from the Air Force Research Laboratory (FA86511010001); G.V. and J.R.M. from the French ANR (FlyBrainImaging/ANR-11-BSV04-003-01); E.B., C.C., R.S. and J.H. from the BBSRC (BB/J017221/1 and BB/J018589/1), and R.D. holds a Wellcome Trust Senior Investigator Award (098362/Z/12/Z).

We thank colleagues listed in the extended methods section of *Supplementary Information*, as well as the Developmental Studies Hybridoma Bank and the Bloomington Stock Center for providing antibodies and fly strains. We are grateful to N.J. Buckley for support during the early stages of this study, to J. Harris from the Nikon-KCL imaging centre for help with super resolution microscopy, and to K.P. Giese, S. Birman, I. Salecker, R. Strauss, J. Simpson, B. van Swinderen, W. Schultz, R.M. Costa, P. Redgrave and T. Prescott for helpful comments and discussions.

### AUTHOR CONTRIBUTIONS

Z.N.L. & D.C.D. performed and analysed anatomical experiments. A.S. performed and analysed GRASP experiments. G.V. & J.R.M. performed and analysed bioluminescence imaging. E.B. & J.J.L.H. performed and analysed electrophysiology and GCaMP imaging experiments. B.K. performed and analysed behavioural experiments. R.F. designed DART software including custom-made MATLAB scripts; J.D. & D.M.H. wrote earlier versions of MATLAB scripts. R.F. & J.D. performed statistical analyses. V.G.F. & R.J.D. designed and analysed computational model; S.B. & B.de B. provided early computational modelling. C.C. & R.S. performed and analysed temperature preference experiments. Y.A., B.X., K.E.W. & D.A.S. performed molecular and anatomical experiments. S.J.S. supervised super resolution microscopy. K.E. & K.I. generated and identified *NV45lexA* line. S.B. & N.J.S. performed and analysed EM experiments, and N.J.S. conducted comparative anatomical analyses. F.H. directed the study and prepared the manuscript to which the authors contributed.

## REFERENCES

1. Strausfeld, N.J. & Hirth, F. Deep homology of arthropod central complex and vertebrate basal ganglia. Science 340, 157-161 (2013).

2. Kravitz, A.V., et al. Regulation of parkinsonian motor behaviours by optogenetic control of basal ganglia circuitry. Nature 466, 622-626 (2010).

3. Cui, G., et al. Concurrent activation of striatal direct and indirect pathways during action initiation. Nature 494, 238-242 (2013).

4. Isomura, Y., et al. Reward-modulated motor information in identified striatum neurons. J. Neurosci. 33, 10209-10220 (2013).

5. Mink, J.W. The basal ganglia: focused selection and inhibition of competing motor programs. Prog. Neurobiol. 50, 381-425 (1996).

6. Graybiel, A.M. The basal ganglia. Curr. Biol. 10, R509-511 (2000).

7. Grillner, S., Hellgren, J., Ménard, A., Saitoh, K., and Wikström, M.A. Mechanisms for selection of basic motor programs – roles for the striatum and pallidum. Trends Neurosci. 28, 364-370 (2005).

8. Nelson, A.B. & Kreitzer, A.C. Reassessing models of basal ganglia function and dysfunction. Annu. Rev. Neurosci. 37, 117-135 (2014).

9. Jin, X. & Costa, R.M. Shaping action sequences in basal ganglia circuits. Curr. Opin. Neurobiol. 33, 188-196 (2015).

10. Fiore, V.G., Dolan, R.J., Strausfeld, N.J., & Hirth, F. Evolutionarily conserved mechanisms for the selection and maintenance of behavioural activity. Philos. Trans. R. Soc. Lond. B. Biol. Sci. 370, 20150053 (2015).

11. Strauss, R. & Heisenberg, M. A higher control center of locomotor behavior in the *Drosophila* brain. J. Neurosci. 5, 1852-1861 (1993).

12. Martin J.R., Raabe T., & Heisenberg, M. Central complex substructures are required for the maintenance of locomotor activity in *Drosophila melanogaster*. J. Comp. Physiol A 185, 277-288 (1999).

13. Martin, J., Faure, P. & Ernst, R. The power law distribution for walking-time intervals correlates with the ellipsoid-body in *Drosophila*. J. Neurogenet. 15, 205-219 (2001).

14. Boll, W. & Noll, M. The *Drosophila Pox neuro* gene: control of male courtship behaviour and fertility by a complete dissection of all enhancers. Development 129, 5667-5681 (2002).

15. Sakura, M., Lambrinos, D., & Labhart, T. Polarized skylight navigation in insects: model and electrophysiology of e-vector coding by neurons in the central complex. J. Neurophysiol. 99, 667-682 (2007).

16. Neuser, K., Triphan, T., Mronz, M., Poeck, B. & Strauss, R. Analysis of a spatial orientation memory in *Drosophila*. Nature 453, 1244-1247 (2008).

17. Pan, Y., et al. Differential roles of the fan-shaped body and the ellipsoid body in *Drosophila* visual pattern memory. Learn. Mem. 5, 289-295 (2009).

18. Lebestky, T., et al. Two different forms of arousal in *Drosophila* are oppositely regulated by the dopamine D1 receptor ortholog DopR via distinct neural circuits. Neuron 64, 522-536 (2009).

19. Heinze, S., Gotthardt, S., & Homberg, U. Transformation of polarized light information in the central complex of the locust. J. Neurosci. 29, 11783-11793 (2009).

20. Kong, E.C., et al. A pair of dopamine neurons target the D1-like dopamine receptor DopR in the central complex to promote ethanol-stimulated locomotion in *Drosophila*. PLoS One 5, e9954 (2010).

21. Ofstad, T.A., Zuker, C.S., & Reiser, M.B. Visual place learning in *Drosophila melanogaster*. Nature 447, 204-207 (2011).

22. Seelig, J.D. & Jayaraman, V. Feature detection and orientation tuning in the Drosophila central complex. Nature 503, 262-266 (2013).

23. Guo, P. & Ritzmann, R.E. Neural activity in the central complex of the cockroach brain is linked to turning behaviors. J. Exp. Biol. 216, 992-1002 (2013).

24. Kathman, N.D., Kesavan, M., & Ritzmann, R.E. Encoding wide-field motion and direction in the central complex of the cockroach Blaberus discoidalis. J. Exp. Biol. 217, 4079-4090 (2014).

25. Martin, J.P., Guo, P., Mu, L., Harley, C.M, & Ritzmann RE. Central-complex control of movement in the freely walking cockroach. Curr. Biol. 25, 2795-2803 (2015).

26. Seelig, J.D., and Jayaraman, V. Neural dynamics for landmark orientation and angular path integration. Nature 521, 186-191 (2015).

27. Liu, S., Liu, Q., Tabuchi, M., & Wu, M.N. Sleep Drive Is Encoded by Neural Plastic Changes in a Dedicated Circuit. Cell 165, 1347-1360 (2016).

28. Owald, D., Lin, S., & Waddell, S. Light, heat, action: neural control of fruit fly behaviour. Philos. Trans. R. Soc. Lond. B. Biol. Sci. 370, 20140211 (2015).

29. Hanesch, U., Fischbach, K.-F., & Heisenberg, M. Neuronal architecture of the central complex in *Drosophila melanogaster*. Cell Tissue Res. 257, 343-366 (1989).

30. Renn, S.C., et al. Genetic analysis of the *Drosophila* ellipsoid body neuropil: organization and development of the central complex. J. Neurobiol. 41, 189-207 (1999).

31. Young, J.M. & Armstrong, J.D. Structure of the adult central complex in Drosophila: organization of distinct neuronal subsets. J. Comp. Neurol. 518, 1500-1524 (2010).

32. Hirth, F., et al. An urbilaterian origin of the tripartite brain: developmental genetic insights from *Drosophila*. Development 130, 2365-2373 (2003).

33. Urbach, R. & Technau, G.M. Neuroblast formation and patterning during early brain development in *Drosophila*. BioEssays 26, 739-751 (2004).

34. Roy, B., et al. Metamorphosis of an identified serotonergic neuron in the *Drosophila* olfactory system. Neural Dev. 2, 20 (2007).

35. Boyan, G. & Liu, Y. Timelines in the insect brain: fates of identified neural stem cells generating the central complex in the grasshopper *Schistocerca gregaria*. Dev. Genes Evol. 224, 37-51 (2014).

36. Nicolaï, L.J., et al. Genetically encoded dendritic marker sheds light on neuronal connectivity in *Drosophila*. Proc. Natl. Acad. Sci. USA 107, 20553-20558 (2010).

37. Feinberg, E.H., et al. GFP Reconstitution Across Synaptic Partners (GRASP) defines cell contacts and synapses in living nervous systems. Neuron 57, 353-363 (2008).

38. Gordon, M.D. & Scott, K. Motor control in a Drosophila taste circuit. Neuron 61, 373-384 (2009).

39. Kahsai, L. & Winther, A.M. Chemical neuroanatomy of the *Drosophila* central complex: distribution of multiple neuropeptides in relation to neurotransmitters. J. Comp. Neurol. 519, 290-315 (2011).

40. Martin, J.R., Rogers, K.L., Chagneau, C., & Brûlet, P. In vivo bioluminescence imaging of Ca^2+^ signalling in the brain of *Drosophila*. PLoS One 2, e275 (2007).

41. Sayeed, O. & Benzer, S. Behavioral genetics of thermosensation and hygrosensation in *Drosophila*. Proc. Natl. Acad. Sci. USA 93, 6079-6084 (1996).

42. Dillon, M.E., Wang, G., Garrity, P.A., & Huey, R.B. Thermal preference in Drosophila. J. Therm. Biol. 34, 109-119 (2009).

43. Hamada, F.N., et al. An internal thermal sensor controlling temperature preference in Drosophila. Nature 454, 217-220 (2008).

44. Tang, X., Platt, M.D., Lagnese, C.M., Leslie, J.R., & Hamada, F.N. Temperature integration at the AC thermosensory neurons in Drosophila. J. Neurosci. 33, 894-901 (2013).

45. Ito, M., Masuda, N., Shinomiya, K., Endo, K. & Ito, K. Systematic analysis of neural projections reveals clonal composition of the Drosophila brain. Curr. Biol. 23, 644-655 (2013).

46. Chen, T.-W. et al. Ultrasensitive fluorescent proteins for imaging neuronal activity. Nature 499, 295-300 (2013).

47. Bhatia, K.P. & Marsden, C.D. The behavioural and motor consequences of focal lesions of the basal ganglia in man. Brain 117, 859-876 (1994).

48. Zeng, L., Leplow, B., Höll, D., & Mehdorn, M. Quantification of human spatial behavior in an open field-locomotor maze. Percept. Mot. Skills. 97, 917-935 (2003).

49. Cisek, P. & Kalaska, J.F. Neural mechanisms for interacting with a world full of action choices. Annu. Rev. Neurosci. 33, 269-298 (2010).

50. Sweeney, S.T., Broadie, K., Keane, J., Niemann, H., & O’Kane, C. J. Targeted expression of tetanus toxin light chain in *Drosophila* specifically eliminates synaptic transmission and causes behavioral defects. Neuron 14, 341-351 (1995).

51. Nitabach, M.N., et al. Electrical hyperexcitation of lateral ventral pacemaker neurons desynchronizes downstream circadian oscillators in the fly circadian circuit and induces multiple behavioral periods. J. Neurosci. 26, 479-489 (2006).

52. Gonzalez-Bellido, P.T., Wardill, T.J., Kostyleva, R., Meinertzhagen, I.A., & Juusola, M. Overexpressing temperature-sensitive dynamin decelerates phototransduction and bundles microtubules in Drosophila photoreceptors. J. Neurosci. 29, 14199-14210 (2009).

53. Diaper, D.C., et al. Loss and gain of Drosophila TDP-43 impair synaptic efficacy and motor control leading to age-related neurodegeneration by loss-of-function phenotypes. Hum. Mol. Genet. 22, 1539-1557 (2013).

54. Coulom, H. & Birman, S. Chronic exposure to rotenone models sporadic Parkinson’s disease in Drosophila melanogaster. J. Neurosci. 24, 10993-10998 (2004).

55. Deco, G., Jirsa, V.K., Robinson, P.A., Breakspear, M., & Friston, K. The dynamic brain: from spiking neurons to neural masses and cortical fields. PLoS Comput. Biol. 4, e1000092 (2008).

56. Lin, C.Y., et al. A comprehensive wiring diagram of the protocerebral bridge for visual information processing in the *Drosophila* brain. Cell Rep. 3, 1739-1753 (2013).

57. Wolff, T., Iyer, N.A., & Rubin, G.M. Neuroarchitecture and neuroanatomy of the Drosophila central complex: A GAL4-based dissection of protocerebral bridge neurons and circuits. J. Comp. Neurol. 523, 997-1037 (2015).

58. Shih, C.T., et al. Connectomics-based analysis of information flow in the Drosophila brain. Curr Biol. 25, 1249-58 (2015).

59. Rabinovich, M.I., Verona, P., Selverston, A.I., & Abarbanel, H.D.I. Dynamical principles in neuroscience. Reviews of Modern Physics 78, 2013-1265 (2006).

60. el Jundi, B., Warrant, E.J., Byrne, M.J., Khaldy, L., Baird, E., Smolka, J. & Dacke, M. Neural coding underlying the cue preference for celestial orientation. Proc. Natl. Acad. Sci. USA 112, 11395-11400 (2015).

61. Tomchik, S.M. & Davis, R.L. Out of sight, but not out of mind. Nature 453, 1192-1194 (2008).

62. Strausfeld, N.J. Arthropod Brains: Evolution, Functional Elegance and Historical Significance (Cambridge MA: Harvard University Press) (2012).

63. Homberg, U. Flight-correlated activity changes in neurons of the lateral accessory lobes in the brain of the locust *Schistocerca gregaria*. J. Comp. Physiol. 175, 597-610 (1994).

64. Namiki, S., et al. Information flow through neural circuits for pheromone orientation. Nat. Commun. 5, 5919 (2014).

65. Jin, X., Tecuapetla, F., & Costa, R.M. Basal ganglia subcircuits distinctively encode the parsing and concatenation of action sequences. Nat. Neurosci. 17, 423-430 (2014).

66. Tecuapetla, F., Matias, S., Dugue, G.P., Mainen, Z.F., & Costa, R.M. Balanced activity in basal ganglia projection pathways is critical for contraversive movements. Nat. Commun. 5, 4315 (2014).

67. Vicente, A.M., Galvão-Ferreira, P., Tecuapetla, F., & Costa, R.M. Direct and indirect dorsolateral striatum pathways reinforce different action strategies. Curr. Biol. 26, R267-269 (2016).

68. Redgrave, P., Prescott, T.J., & Gurney, K. The basal ganglia: a vertebrate solution to the selection problem? Neuroscience 89, 1009-1023 (1999).

69. Gunaydin, L.A. & Kreitzer, A.C. Cortico-basal ganglia circuit function in psychiatric disease. Annu. Rev. Physiol. 78, 327-350 (2016).

70. Faville, R., Kottler, B., Goodhill, G.J., Shaw. P.J., & van Swinderen, B. How deeply does your mutant sleep? Probing arousal to better understand sleep defects in *Drosophila*. Sci Rep. 5, 8454 (2015).

